# A representation of the visual environment in the human basolateral amygdala supports learning and choice

**DOI:** 10.64898/2026.10.07.757347

**Authors:** Grace Jang, Kierstin Riels, Elias Aberman, Grace Tao, Philip A. Kragel

**Affiliations:** Neuroscience Graduate Program, Emory University, Atlanta, Georgia, USA; Department of Psychology, Emory University, Atlanta, Georgia, USA; Department of Psychiatry and Behavioral Sciences, Emory University, Atlanta, Georgia, USA

## Abstract

Animals continuously transform sensory inputs into representations that guide behavior. Although the amygdala is a brain structure critical for learning and choice, how natural sensory experience is represented within this circuit remains unknown. Here, we identify a representation of the visual environment in the human basolateral amygdala (BLA) that contains information relevant for learning and choice. Using image-computable encoding models trained on responses to naturalistic videos, we recover a continuous, low-dimensional representation of visual experience in the BLA. The representation generalizes across individuals and independent datasets and prospectively predicts BLA responses to novel visual stimuli generated through closed-loop image synthesis. Signals derived from the representation also informed reinforcement-learning models of human choice. Together, these findings show that the human BLA organizes complex visual experience into a structured representation that can support learning and flexible behavioral responses.

---

A central challenge for any intelligent system—biological or artificial—is how to transform high-dimensional, naturalistic sensory inputs into low-dimensional representations that drive flexible, state-dependent behavior. Natural environments contain more sensory variation than an organism can represent explicitly at any moment. Animals must extract regularities that remain useful as they encounter new objects, scenes, and events. Ecological theories of perception emphasize this relationship between environmental structure and behavior^1^. Reinforcement-learning theories describe how experience can update future choices^2^, but they do not specify how complex sensory inputs are organized into neural states over which this learning can operate. How the brain transforms natural sensory experience into representations that can inform choice remains unknown.

Intracranial self-stimulation provided an early demonstration that activity in subcortical circuits can reinforce behavior^3,4^. Subsequent electrical stimulation studies in nonhuman animals showed that activation of these circuits can support learned behavior and engage neural signals associated with reinforcement^5,6^. More recent optogenetic studies have shown that activation of defined subcortical populations can similarly support conditioning^7–10^. These manipulations establish that activity in these circuits can influence learning, but they bypass the sensory pathways through which organisms normally encounter the environment. How naturally occurring sensory experiences are transformed into neural signals that can guide learning and choice remains unknown.

The basolateral amygdala (BLA) is well positioned to link visual experience with behavioral consequences^11^. It receives high-level visual information from the ventral temporal cortex and converges with signals related to physiological states and behavioral outcomes ^12,13^. Consistent with this organization, recordings and functional imaging in nonhuman primates demonstrate that the BLA responds selectively to biologically significant visual objects ^14,15^, including faces, animals, food, and threats^14–17^, as well as unconditioned appetitive and aversive stimuli beyond the visual domain^10,18–21^. The amygdala also contributes to reinforcement learning and choice^13,22–24^, raising the possibility that this region transforms complex sensory inputs into a lower-dimensional representation that provides structured states over which behavioral significance can be learned^25^.

Recent work has made it possible to study this sensory organization directly. Image-computable models of the ventral visual stream predict BLA responses in nonhuman primates^15,18,26^ and humans^27^. In nonhuman primates, these models predict population activity that links natural visual inputs with the representation of unconditioned gustatory and tactile valence^18^. Population recordings further show that BLA activity can form continuous, low-dimensional representations of emotional states^28^. Together, these findings indicate that BLA activity contains structured information about both sensory experience and behavioral state. They do not establish how natural visual experience is organized within the BLA or whether that organization contains information that can be used for learning.

Here, we combine naturalistic fMRI, image-computable encoding models, deep image synthesis, and reinforcement-learning models to address these questions. We first recover a representation of the visual environment from BLA responses during naturalistic movie viewing and test whether it generalizes across individuals and independent datasets. We then use the recovered representation to generate novel visual stimuli and test whether it predicts BLA responses to those stimuli. Finally, we ask whether information in the representation could account for trial-by-trial changes in choice behavior. Together, these experiments test whether a representation recovered from natural visual experience can generalize beyond the experiences used to estimate it, predict responses to novel inputs, and provide structured information that can be incorporated into learning.

## Characterizing a representation of the visual environment in human BLA

Recent work demonstrates that primate BLA responses can be predicted by image-computable models of the visual system^15,16^. We therefore asked whether natural visual experience could be used to recover a shared representation of the visual environment in the human BLA.

We fit voxel-wise encoding models that used a high-dimensional feature space to predict BOLD signal in the BLA and inferior temporal cortex (IT) acquired while participants viewed a naturalistic, emotional film. We then used these models to synthesize images predicted to maximize activity in each region. In a cross-model validation test, BLA models predicted larger responses to BLA-optimized (*M*_BLA_ = 5.452, *SD*_BLA_ = 3.059) than IT-optimized images (*M*_IT_ = 3.311, *SD*_IT_ = 2.438; *t*_37_ = 6.059, *p* < .001, 95% CI = [1.425, 2.856]), whereas IT models exhibited complementary selectivity (*M*_IT_ = 2.15, *SD*_IT_ = 1.103, *M*_BLA_ = 1.081, *SD*_BLA_ = 1.111, t_37_ = 13.778, *p* < .001, 95% CI = [0.912, 1.226]; Fig. 1a). Thus, models estimated from naturalistic visual experience recovered distinct BLA– and IT-selective representations that could be interrogated with novel images.

**Figure 1.**
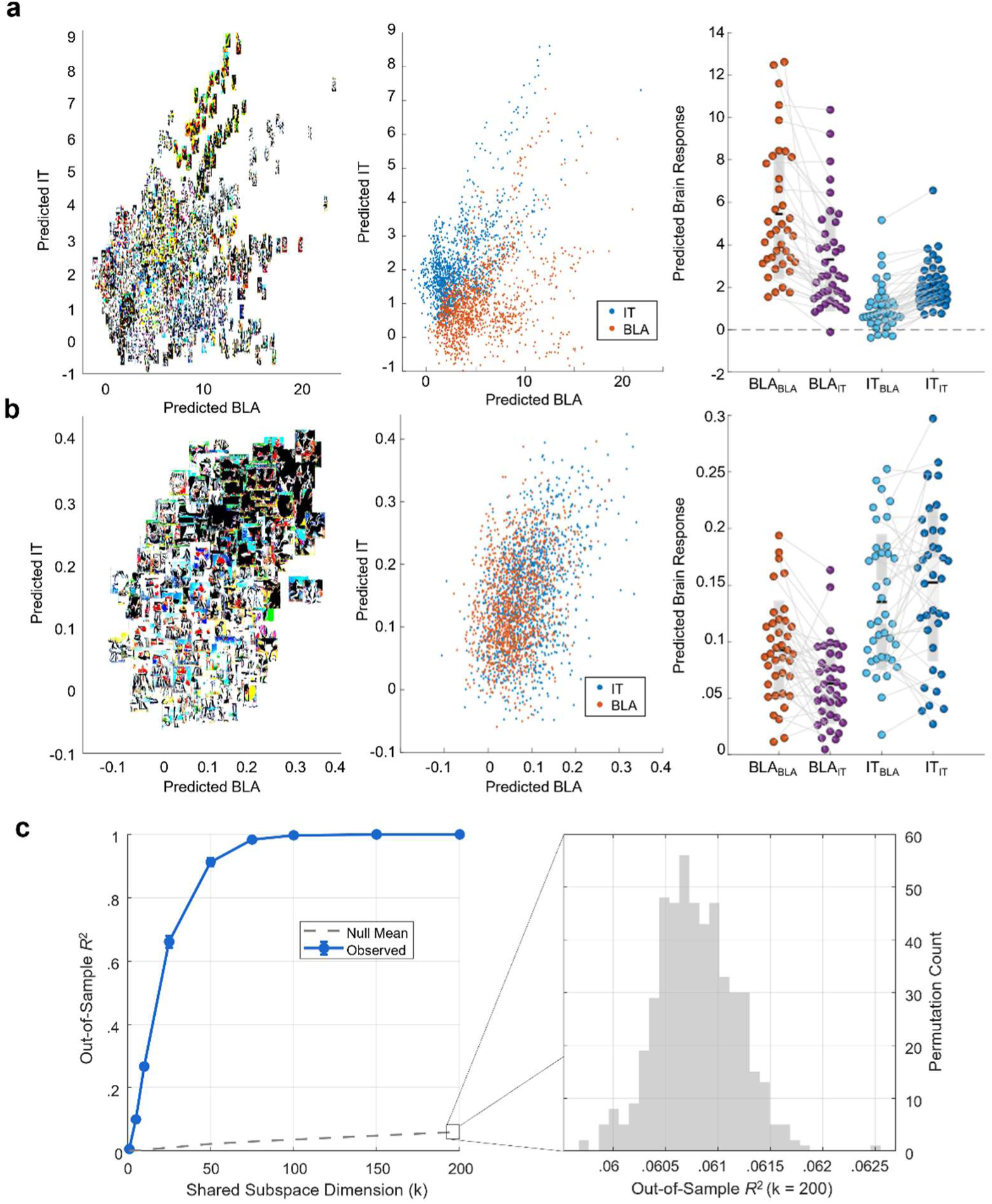
Recovering and validating BLA-selective visual representations. **a**, Model validation using individualized encoding models fit during naturalistic movie viewing. Synthetic visual images generated show relationships between predicted responses and low-level visual features. Scatter plots (middle) illustrate predicted IT vs. BLA model responses for IT-(blue) and BLA-optimized (orange) images. Group-level comparisons (right) demonstrate cross-model selectivity, showing higher predicted responses in targeted regions. **b,** Generalization of an independent population-level encoding model. Synthetic image distribution (left) and predicted response scatter plots (middle) show model performance on novel synthetic stimuli. Model predictions (right) demonstrate that BLA-optimized stimuli consistently evoke higher predicted BLA responses across independent datasets and participants. **c,** Cross-subject alignment of the mapping between visual features and amygdala BOLD response patterns for subspaces up to 200 dimensions. Blue line indicates the performance of leave-one-subject-out cross validation. Gray histogram shows the randomization null at *k* = 200 dimensions.

To test whether the encoding models captured a functional geometry shared across individuals, we evaluated the geometry of model coefficients using leave-one-subject-out cross-validation. For each held-out participant, a shared basis was estimated from the remaining 37 participants and used to reconstruct the held-out encoding model coefficients. The shared representation captured 98% of out-of-sample variance with 75 dimensions and 99.8% with 100 dimensions (Fig. 2c), demonstrating that independently estimated encoding models shared a highly conserved, low-dimensional structure within the 4,096-dimensional feature space.

**Figure 2.**
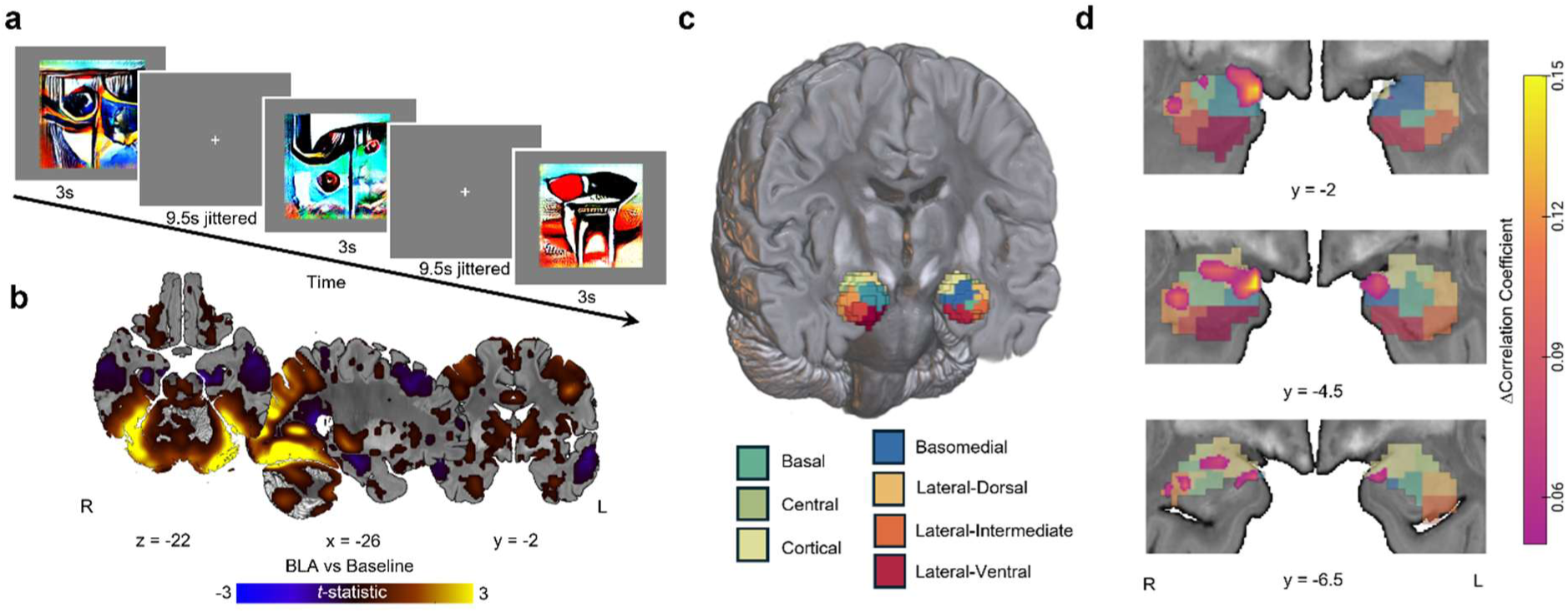
Prospective validation of the BLA manifold. **a**, Directed viewing experimental paradigm. Participants viewed synthetic images for 3 seconds each, interleaved with a jittered fixation cross (mean inter-trial interval = 9.5 s), while performing a change-detection task. **b,** Whole-brain fMRI response to synthetic BLA-optimized stimuli versus baseline. Viewing BLA-optimized images recruited widespread activation across the ventral visual stream, amygdala, and connected subcortical and prefrontal networks. **c,** Volumetric rendering showing segmentation of the human amygdala into distinct anatomical subdivisions. **d,** Voxel-wise encoding model comparisons within the amygdala across coronal slices. Amygdala BOLD signal variations were significantly better predicted by BLA-derived encoding models than IT-derived models, localized primarily within basolateral subregions.

Having established a shared structure across individuals, we next asked whether the representation generalized to independent datasets and novel experimental conditions. We evaluated an independent, population-level BLA encoding model estimated from a separate cohort viewing a different feature film^27^. Despite differences in participants, stimuli, and acquisition parameters, this model predicted larger responses to BLA-optimized (*M* = 0.091, *SD* = 0.042) than IT-optimized images (*M* = 0.063, *SD* = 0.034; t_37_ = 4.067, 95% CI = [.0145, 0417], *p* < .001; Fig 2b).

We next characterized the visual information captured by the recovered representation. We analyzed the optimized images using artificial neural networks for visual salience^29^ and deep convolutional neural networks designed to detect higher-level semantic^30^ and affective^31^ content (Supplementary Fig. S1). Visual salience was modestly related to predicted BLA responses (*r* = .1663 ± .0370 (sem), 95% CI = [.0976, .2414]). In contrast, object and emotion category predictions were largely unrelated to one another (*r* = .0371 ± .0031 (sem), 95% CI = [.032 .044]), and each showed only weak relationships with predicted BLA responses (*r*_emotion_= .0606 ± .0531 (sem), 95% CI = [-.0541 .1604]; *r*_object_= .106 ± .0407 (sem), 95% CI = [.0229 .1846]; Supplementary Tables 1-2). Direct comparison of low-level visual properties confirmed that BLA– and IT-optimized images were similar in terms of global spatial contrast and color saturation (Supplementary Fig. S2). These findings indicate that the representation was not well characterized by individual semantic or affective categories. Instead, it reflected combinations of visual features distributed across the natural environment.

## Prospectively testing BLA representations with novel visual inputs

Having identified a representation of the visual environment in the human BLA, we next asked whether it could predict neural responses to visual inputs that were not present during model estimation. We generated synthetic images from both individualized and population-level encoding models and presented them during fMRI while participants performed a simple change-detection task.

Participants successfully discriminated between successive stimuli, indicating that they engaged with the images (*n* = 35; *d*^’^ = .4910 ± .1068 (sem), 95% CI = [.2576, .6322] for population stimuli; *d*^’^ = .4244 ± .1076 (sem), 95% CI = [.2053, .6322] for individualized stimuli). Post-scan ratings of emotional experience suggested the stimuli did not produce large changes in subjective emotion or affect (Supplementary Tables 3-4). Both BLA– and IT-optimized stimuli evoked robust responses in the ventral visual stream, the amygdala, and multiple areas with known connectivity to the BLA, including the dorsomedial prefrontal cortex^32^, mediodorsal thalamus^33^, and striatum^34^ (Fig. 2b).

We then tested the model predictions for each synthetic image. Predicted BLA responses were systematically related to observed trial-by-trial variation in amygdala activity, and BLA-derived predictions explained amygdala responses better than IT-derived predictions (peak *t* = 5.27, MNI_x,y,z_ = [24, –3, –19], extent = 8 voxels; Fig. 2d). Thus, a representation recovered from naturalistic visual experience predicted BLA responses to novel visual inputs.

## BLA representations of the visual environment inform learning and choice

A representation of the visual environment is behaviorally relevant if information about its states can be incorporated into the computations that update future choices. We therefore asked whether BLA-derived information could contribute to learning, and whether the behavioral significance assigned to represented states was shared across participants or varied with individual learning history.

Participants learned cue–outcome contingencies through trial and error, with choices yielding individualized BLA-optimized images, IT-optimized images, or no visual outcome (Fig. 3a; *n* = 39). We incorporated the predicted BLA responses to each outcome image into a *Q*-learning model, allowing the representation to provide trial-specific signals for updating action values (Fig. 3b; see Supplementary Tables 5–7 for the reward structure, parameter recovery, and model recovery). We compared models in which visual outcomes were assigned fixed positive or negative values with models in which the learning signal was determined by predicted BLA responses.

**Figure 3.**
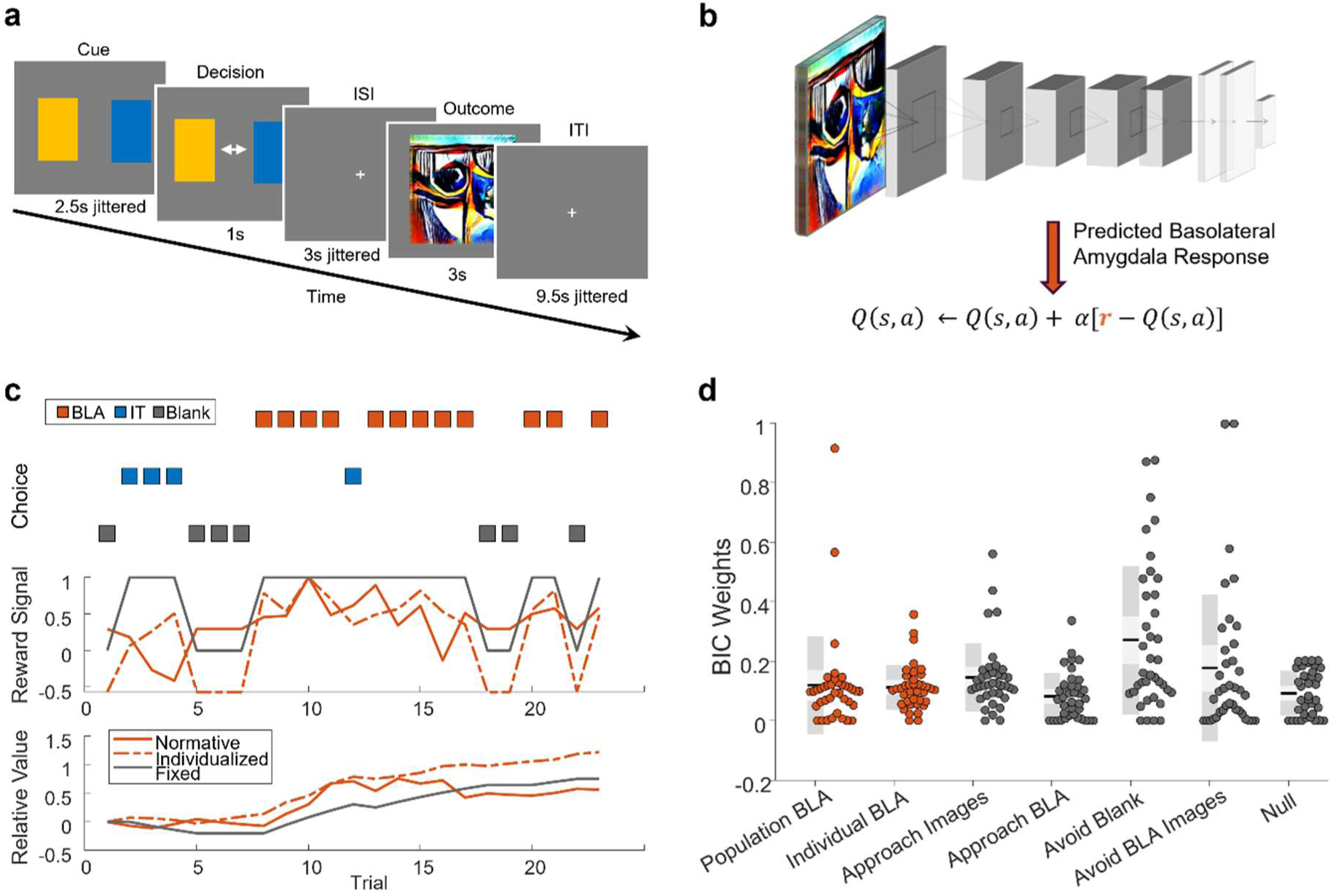
BLA representations of the visual environment can influence learning and choice. **a**, Experimental paradigm. Participants chose between colored cues to receive outcomes consisting of individualized BLA-optimized images, IT-optimized images, or no visual outcome (blank fixation). **b,** *Q*-learning framework. Predicted BLA responses generated by passing outcome images through the encoding model served as dynamic reward teaching signals to update action values via prediction errors. **c,** Representative single-subject learning dynamics across trials. Top panel displays one participant’s choices (associated with BLA, IT, or blank outcomes). Middle and bottom panels illustrate dynamic trial-by-trial reward signals and estimated relative values modeled using population BLA, individualized BLA, and binary fixed-reward assumptions. **d,** BIC weights across participants for the seven candidate models. Continuous BLA models and fixed-value accounts received support, while models assigning negative value to BLA-optimized images were strongly favored in a subset of participants, revealing individual differences in the behavioral value of the synthesized images.

At the population level, choices were most consistently explained by learning to avoid the absence of a visual outcome. The model assigning a negative value to blank trials had the largest support (mean *w*_BIC_ = .2707, 95% CI = [.2013 .3581]; expected probability = 0.532), and models using image-computable rewards received partial support (mean *w*_BIC_= .1159, 95% CI = [.0953 .1531], expected probability = 0.130; see Supplementary Table 8).

Individual fits revealed substantial heterogeneity in how participants assigned behavioral significance to the synthesized images. This variation was evident despite the shared BLA representational structure. When isolating the best-fitting model for each participant (ΔBIC = 0), avoidance-driven strategies best accounted for 71.05% (27/38) of the sample. In contrast, models in which images had positive or zero value accounted for 23.68% (9/38) of participants, whereas the models in which reward was derived from predicted BLA activity provided the best fit for only 5.26% (2/38). Despite this variation, one of the two BLA-derived learning models still provided a competitive account of choice (ΔBIC < 2) for 71.05% of the sample (27/38). In sum, many participants were well described using fixed outcome values and by learning signals that varied with the predicted BLA response. These results suggest that information in the shared representation can be incorporated into learning without requiring that its sensory states acquire a common behavioral value.

## Discussion

How the amygdala organizes sensory information that can support different behaviors remains an open question. On the one hand, research has revealed functionally selective circuits within the amygdala^35–38^, including BLA populations associated with opposing valence and behavioral outcomes^39,40^. On the other hand, considerable evidence suggests that BLA population activity can vary with physiological state and behavioral context^41^. Population recordings have nevertheless revealed multidimensional representations that combine sensory, affective, and behavioral information^42–45^. Together, this evidence suggests that circuit specialization and flexible population representations coexist within the amygdala.

The present findings clarify this duality by identifying a continuous organization of the visual environment in the human BLA that generalizes across individuals and experiences and predicts responses to novel stimuli. Despite this shared representational structure, the behavioral consequences assigned to represented states varied across participants. At the population level, choices were most consistently explained by learning to avoid the absence of visual stimulation, whereas models incorporating BLA-derived learning signals provided competitive accounts for a substantial subset of participants. Thus, the BLA representation could contribute to learning without imposing a common behavioral meaning on its states^46^.

This distinction is important for understanding the relationship between sensory representation and valuation. The amygdala has long been implicated in learning the behavioral significance of stimuli^47,48^ and work in primates has shown that value representations in the amygdala can change rapidly when reinforcement contingencies are reversed^17,20,49^. Genetically defined BLA populations can also preferentially support positively and negatively valenced behaviors and acquire predictive responses through learning^40^. Such findings demonstrate that amygdala value signals are state– and experience-dependent. If reinforcement learning operates over sensory representations^50^, those representations constrain the space over which value can be learned^51^. Stimuli that occupy nearby states in the representation can be treated as related, whereas differences represented along other dimensions can support distinct learning. Our results identify such a structured representation in the BLA.

This perspective builds on recent work linking visual inputs to amygdala responses across species. Image-computable models have related natural visual features to BLA responses in nonhuman primates^14,17^, while population recordings in rodents have revealed continuous geometries of amygdala activity^28^. In humans, naturalistic encoding models similarly relate visual input to amygdala responses and affective dimensions^27^. The present findings extend this work by showing that a recovered representation can predict BLA responses to stimuli generated after model estimation and that information in this representation can contribute to subsequent learning and choice.

These findings link a representation of the sensory environment to reinforcement learning in humans. Intracranial self-stimulation demonstrated that electrical stimulation of the brain can drive reinforcement^3,4^. Subsequent optogenetic studies have shown that activation of defined subcortical populations can support conditioning and reinforce specific behavioral responses^5–7^. These manipulations, however, bypass the sensory computations through which organisms naturally encounter the world. Our results identify a representation of the visual environment that can participate in reinforcement learning. The BLA can link sensory states to the computations that guide behavior^52^, providing a neural substrate through which humans can learn the behavioral consequences of the world they encounter.

The organization of this representation may help explain how the behavioral significance of sensory states can remain flexible while the structure of the state space remains stable. The shared geometry was sufficiently conserved to generalize across individuals and naturalistic experiences, while the behavioral consequences associated with states represented in the BLA varied across participants. Such organization may arise through interactions between higher-level visual cortex and the BLA^53^, together with recurrent influences from other cortical and subcortical systems^54^. Future research using comparative and developmental methods is needed to determine which aspects of this organization are conserved and which emerge through experience. In particular, longitudinal studies could reveal how its geometry changes as sensory states acquire new behavioral consequences^55^.

More broadly, our findings highlight that the utility of a theory of amygdala function depends on the level at which it explains behavior. Value and affect describe important properties of the consequences that guide learning, but they do not specify how sensory states are organized so that those consequences can be learned. A representation of the environment specifies the states over which learning can operate and thereby constrains the behavioral significance that those states can acquire. The human BLA contains such a representation—one that links the visual environment to learning and choice while leaving the behavioral significance of its states flexible. Understanding how this representational structure is acquired, conserved, and reshaped by experience may be central to understanding how the amygdala supports adaptive behavior.

## Methods

Given evidence that the amygdala encodes the valence afforded by the sensory environment and is involved in learning, we designed an experimental pipeline to recover, validate, and functionally evaluate individualized image-computable representations of human BLA activity. All experiments were conducted during a single study visit. In the first fMRI session, participants viewed an emotional movie while we estimated individualized encoding models relating visual features to BLA activity. These models were then used to generate synthetic images predicted to engage either the BLA or inferior temporal cortex (IT) as a control. In the second fMRI session, participants viewed both individualized synthetic images and an independent set of synthetic images generated from a separate cohort^15^ to evaluate whether the recovered representations prospectively predicted BLA responses and generalized across individuals. Subsequently, participants performed a reinforcement learning task in which individualized synthetic images served as learning outcomes. Finally, participants completed a post-scan behavioral task in which they rated their experience of the synthetic images using visual analog scales. Together, these experiments evaluated whether recovered BLA representations could be identified from naturalistic brain responses, prospectively engaged through synthetic sensory stimuli, and support adaptive learning.

### Participants

We recruited 50 healthy adult subjects from the Greater Atlanta Metropolitan Area in Georgia (34 Female, *M*_age_ = 33.67 years, *<u>SD</u>*_age_ = 11.28 years). All participants provided written informed consent prior to participation, and procedures were approved by the Institutional Review Board at Emory University. Participants were eligible if they were between 18 and 65 years of age and able to provide informed consent. Exclusion criteria included a history of epilepsy, head trauma, substance abuse or dependence within the 6 months preceding enrollment, pregnancy (self-reported), current drug use (self-reported), or MRI contraindications. Eight participants did not complete the protocol due to technical issues. Data from two participants were excluded from in-scanner behavioral analyses (directed viewing and reinforcement-learning tasks) due to timing issues during scanning. Subsequent quality-control procedures identified one additional participant with substantial motion artifacts during the movie-viewing scan, resulting in exclusion from further analyses. The final sample size varied by analysis and is reported for each experiment below.

### Experimental setup

All experimental tasks (movie viewing, directed viewing, reinforcement learning, and self-report ratings) were implemented using PsychoPy Version 2023.1.2. Visual stimuli were presented via a projector located behind the scanner. During the reinforcement-learning task, participants indicated their choices using their right index and middle fingers on an MR-compatible button box (Current Designs). Outside the scanner, participants rated individualized synthetic stimuli using a mouse-controlled visual analog scale.

### MRI data acquisition

MRI data were acquired using a 3T Siemens MAGNETOM Prisma Fit MRI scanner (Siemens Healthineers, Erlangen, Germany) at Emory University (Atlanta, GA, USA). Structural images were acquired using a T1-weighted MPRAGE sequence (TR = 1900 ms, TE = 2.26 ms, FOV = 256 mm, GRAPPA acceleration factor = 2, flip angle = 9°, 256 × 256 matrix, 1 mm isotropic voxels, 192 slices, anterior–posterior phase encoding direction) and were used for spatial normalization to Montreal Neurological Institute space. Functional images acquired during the movie-viewing and directed-viewing tasks used a multiband echo-planar imaging sequence developed by the Center for Magnetic Resonance Research, University of Minnesota (R2016; TR = 492 ms, TE = 30.4 ms, FOV = 220 mm, multiband acceleration factor = 8, flip angle = 44°, 82 × 82 matrix, 2.7 mm isotropic voxels, 56 interleaved ascending slices, posterior–anterior phase encoding direction). A pair of fieldmap scans matched to the functional acquisition geometry was collected for susceptibility distortion correction (TR = 7220 ms, TE = 73 ms, flip angle = 90°, refocus flip angle = 180°, phase encoding directions = posterior–anterior/anterior–posterior).

### Processing workflow for fMRI experiments

We implemented a real-time data streaming and processing that enabled estimation of individualized encoding models and generation of synthetic stimuli within a single study visit. During the movie-viewing task, DICOM images were streamed from the MRI scanner, converted to NIfTI format, and subjected to lightweight preprocessing. These data were used to estimate region-specific encoding models, and the resulting model parameters were used to generate synthetic stimuli for subsequent fMRI and behavioral experiments.

### Online MRI preprocessing

To speed up online preprocessing, we undersampled fMRI data acquired during the movie viewing task, retaining every fourth data acquisition (effective TR = 1.968s). Using the SPM12 software, we estimated rigid body motion and performed spatial realignment (quality = 0.9, separation = 4 mm, 5 mm FWHM smoothing kernel for estimation). Functional volumes were then normalized to MNI space using a 12-parameter affine transformation followed by nonlinear deformation to the SPM EPI template^56–58^.

### Encoding model development

Encoding models were fit to predict BOLD signals acquired during the movie viewing task. We used EmoNet, a deep convolutional neural network trained to classify images according to emotional content^31^, as a feature extractor. We extracted activations from the penultimate (fc7) layer, which shares the AlexNet architecture and provides a 4096-dimensional visual feature representation. We temporally convolved activation time series with the canonical double gamma response function implemented in SPM^59^. Functional mappings between model features and multivariate BOLD signals were estimated separately for each region of interest using partial least squares regression (PLS^60^; 20 latent dimensions; MATLAB R2024a `plsregress` function^61^). Separate encoding models were fit for the basolateral amygdala (defined anatomically^62^) and inferior temporal cortex (defined using a multimodal parcellation of the cortex^63^).

### Deep image synthesis

Synthetic images were generated to maximize predicted activity in the basolateral amygdala or inferior temporal cortex using the deep image synthesis framework developed by Nguyen and colleagues^64^. A deep generator network trained on ImageNet was used to generate images optimized within the EmoNet feature space. For each image, optimization began from a random latent code and seed image. The latent representation was iteratively modified using the Sphere Covariance Matrix Adaptation optimizer^65^ to maximize the predicted response obtained from the dot product between EmoNet layer fc7 activations and ROI-specific PLS regression coefficients. Optimization proceeded over 300 iterations for each image. For each target region, 34 artificial stimuli were generated, yielding 68 individualized artificial stimuli per participant.

### Image properties

To assess object– and affect-related visual features, we used deep convolutional networks for object^30^ and emotion^31^ classification. We resized each video frame to 227 × 227 px and obtained model predictions for each of 1,000 object and 20 emotion categories. We also modeled the visual salience of stimuli using a neural network model of visual attention that estimates a priority map of the environment^29^ implemented in the GBVS toolbox^66^.

To quantify whether BLA– and IT-optimized stimulus sets differed in fundamental visual properties, we evaluated low-level image statistics and visual salience across all synthesized images. Following standard image-matching parameters^67,68^ we extracted pixel-wise mean luminance, global spatial contrast (luminance standard deviation), mean HSV saturation, and mean HSV value (brightness) for each image. Distributional differences between BLA– and IT-optimized images were assessed via visual inspection and using Cohen’s *d*.

### Experimental Tasks

#### Movie Viewing Task

In the movie viewing task, participants viewed Chatter^69^, a short emotional film selected because it reliably evokes amygdala responses and has been extensively characterized in the Emo-FILM database. Normative ratings of the video indicate that it tends to elicit a sense of alertness and fear in most viewers^69^. The task began with the presentation of a fixation cross for 8 seconds, followed by the movie, which lasted 405 seconds, and ended with a fixation cross for 10 seconds, yielding a run length of 423 seconds. Participants were instructed to watch the movie and to try not to move and were informed in advance that the movie contained strong emotional content.

#### Directed Viewing Task

In the directed viewing task, participants viewed a sequence of artificial images generated to activate different regions of interest. Participants were instructed to watch the images carefully and indicate via button press (using the right index finger) if the current image appeared to be different from the previous image shown. Six participants were not instructed to perform this task and passively viewed images. Participants completed two runs of this task. One used synthetic images generated from an independent cohort^15,70^, and the other used images generated from each participant’s movie-viewing data.

In each run, participants were presented with a fixation cross for 8 seconds, followed by a series of synthetic images interleaved with fixation crosses during inter-trial intervals (ITI). Each synthetic image was shown for 3 seconds. The ITI was determined through random sampling from a Poisson distribution with λ = 9.5 seconds. There were a total of 36 trials, organized in 4 blocks of 9 trials each. Independent images targeted the BLA, centromedial amygdala, whole amygdala, or inferior temporal cortex. Individualized images targeted only the BLA and inferior temporal cortex. Blocks of trials alternated between different types of stimuli in an order counterbalanced across subjects. Each run ended with the presentation of a fixation cross for 10 seconds.

#### Reinforcement Learning Task

Participants completed a reinforcement-learning task in which they learned cue–outcome associations through trial and error. Outcomes consisted of individualized BLA-optimized synthetic images, individualized IT-optimized synthetic images, or no visual outcome. The task comprised three blocks of eight trials. Two blocks assessed learning relative to no visual outcome (BLA versus no outcome; IT versus no outcome), and a third block assessed choices between BLA– and IT-optimized outcomes.

On each trial, participants were presented with a pair of cues and selected one using either the index or middle finger of their right hand (to choose left or right, respectively). The response period lasted 2.5 seconds. Once participants made their response, the selected cue was highlighted for the remainder of the response period, followed by a jittered inter-stimulus interval (mean 3 s; Poisson distribution with λ = 2 s plus a 1 s minimum). The selected outcome was then presented for 2 s as either an individualized synthetic image or a fixation cross (no-outcome condition), followed by a jittered intertrial interval (λ = 9.5 s). Outcome contingencies remained fixed within each block.

#### Self-report Task

After scanning, participants completed an experiential rating task in which they viewed the individualized synthetic stimuli presented during scanning and rated their subjective experience. Each of the 68 individualized synthetic images (34 BLA-optimized and 34 IT-optimized) was presented for 2 s, followed by two rating screens. Stimuli were presented in eight blocks (8–9 trials per block) that alternated between BLA– and IT-optimized images.

Participants rated valence, arousal, and the extent to which each image evoked seven emotion categories (happy, fear, sadness, surprise, anger, disgust, and neutral) using visual analog scales. Valence and arousal ratings were collected during a 6 s response period, followed by a 12 s response period for the seven emotion ratings. Responses were made using a computer mouse and recorded on continuous visual analog scales with labeled endpoints.

### Offline Preprocessing

Functional MRI data were preprocessed using fMRIPrep 23.1.4^71^, based on Nipype 1.9.2^72^. Anatomical preprocessing included correction for intensity nonuniformity, skull stripping, tissue segmentation, and nonlinear normalization to MNI152NLin2009cAsym space using ANTs^73–77^. Functional preprocessing included susceptibility distortion correction using paired fieldmaps^78^, head-motion correction^79^, co-registration to the anatomical image^80,81^, and spatial normalization. Additional details are available in the fMRIPrep documentation.

### fMRI Analyses

#### GLM Analyses

Following preprocessing, the first 10 data acquisitions for each run were discarded. For the directed viewing task, first-level general linear models (GLMs) were specified and estimated in SPM12^82,83^ with separate regressors for each stimulus condition and 24 motion covariates. For runs using synthetic images generated from an independent dataset, conditions of interest included BLA– and IT-optimized images (*n* = 38 independent participants). Contrast images comparing each condition with implicit baseline were entered into linear mixed effects models (*Y* ∼ Region × Source + (1 + Region | subject)) using CanlabCore Tools^84^. Whole-brain statistical maps were thresholded at a level of *q*_FDR_ < .05. As an exploratory analysis, we compared responses to BLA– and IT-optimized images within the cytoarchitectonically defined amygdala^62^ using linear constrasts (*p* < .005, uncorrected for visualization).

To characterize trial-to-trial correspondence between model predictions and neural responses, we estimated single-trial beta images for each viewed synthetic stimulus. For each stimulus, predicted responses were generated using participant-specific BLA and inferior temporal cortex encoding models by computing the dot product of image features and encoding model weights. Trial-specific BOLD responses were then regressed against BLA and IT model predictions to estimate voxel-wise associations between predicted and observed activity. To directly compare whether amygdala responses were better explained by BLA-versus IT-based predictions, we fit linear mixed-effects models with encoding model identity (BLA versus IT), model source (individualized versus normative), and their interaction as fixed effects, and included subject-specific random intercepts and random slopes for encoding model identity (*Y* ∼ Region × Source + (1 + Region | subject)). For analyses including synthetic images generated from an independent dataset, encoding model source (individualized versus normative) was included as an additional factor to evaluate whether prediction accuracy differed as a function of model source (*p* < .005, uncorrected within the amygdala for visualization).

#### Encoding model evaluation

We evaluated the performance of encoding models by computing voxel-wise correlations between encoding model predictions and observed BOLD activation for each target region in each subject. We used 5-fold cross-validation with random partitioning to estimate model generalization performance^85^. Group-level inference was performed using *t*-tests on Fisher-transformed correlation coefficients (*q*_FDR_ < .05, *n* = 38).

#### Shared Manifold Estimation and Cross-Subject Validation

To evaluate whether a lower-dimensional subspace captures manifold alignment across subjects, we performed leave-one-subject-out cross-validation based on principles of functional alignment ^86^. The dataset comprised a tensor of PLS regression coefficients *B* ∈ ℝ^S^ ^×^ ^F^ ^×^ ^V^, where *S* = 38 subjects, *F* = 4,096 features, and *V* = 847 voxels. For each cross-validation fold *s* ∈ {1, …, *S*}, data from the *s*^th^ subject was extracted and mean-centered along the spatial dimension to form the held-out test matrix 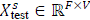. Data from the remaining *S* − 1 training subjects were similarly centered and concatenated to form the group training matrix 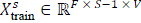. Singular Value Decomposition was computed on 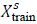 to extract an orthonormal spatial basis *U*^s^ ∈ ℝ^F×F^.

Out-of-sample reconstruction performance was assessed by projecting the mean-centered test data 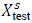 onto the leading k columns of the shared basis 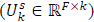:

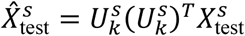

Generalization accuracy was quantified using the out-of-sample proportion of variance explained:

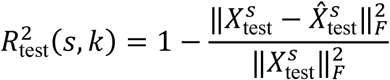

This procedure was systematically evaluated across a range of subspace dimensions k ∈ {1, 5, 10, 25, 50, 75, 100, 150, 200}.

Statistical significance of the out-of-sample variance explained was estimated using a non-parametric feature-permutation test ^87^. To construct the null distribution under the hypothesis of no cross-subject alignment, feature labels were independently scrambled for each test subject:

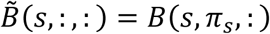

where *π*_s_ represents a unique random permutation vector generated independently for subject s. To prevent data leakage and properly reflect sampling variability under the null hypothesis, each permutation iteration underwent the complete *S*-fold cross-validation pipeline on 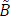. For each permuted draw, a null basis 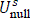 was trained on *S* − 1 permuted subjects and evaluated exclusively on the held-out *S*^th^ permuted subject matrix 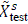. The empirical *p*-value at each dimension *k* was calculated as the proportion of null iterations whose mean out-of-sample *R*^2^ equaled or exceeded the observed mean cross-validated *R*^2^.

#### Encoding model generalization

To evaluate encoding-model selectivity, predicted responses to individualized synthetic images were computed using both the BLA and IT encoding models. Responses were averaged separately for each stimulus type and target model, yielding four conditions (BLA images in BLA models, BLA images in IT models, IT images in IT models, and IT images in BLA models). The same procedure was repeated using synthetic images generated from an independent cohort to assess generalization across participants and datasets.

### Behavioral analyses

#### Reinforcement learning

To determine whether synthetic stimuli that engage the BLA influence adaptive choice behavior, we fit reinforcement-learning models to participants’ choices during the learning task (*n* = 39). We evaluated six competing models describing how participants might assign value to visual outcomes: 1) BLA– and IT-optimized stimuli as equally rewarding with no value assigned to blank outcomes, 2) avoidance of blank outcomes, 3) approach toward BLA-optimized stimuli, 4) avoidance of BLA-optimized stimuli, 5) teaching signals determined by predicted BLA responses from a normative encoding model derived from an independent sample ^15^ and 6) teaching signals determined by predicted BLA responses from individualized encoding models. BLA predictions for Models 5 (normative, group-level scaling) and 6 (individualized scaling) were normalized using a zero-preserving global cohort mean scaling prior to model fitting.

For each model, trial-by-trial action values were estimated using a Rescorla–Wagner *Q*-learning framework. Action values were initialized at zero and updated following each outcome according to a prediction-error learning rule:

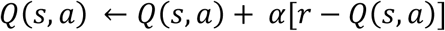

where *α* represents the learning rate and *r* represents the model-specific outcome value. Choices were generated using a softmax function with a temperature parameter β controlling choice stochasticity.

Subject-specific learning parameters (*α* and β) were estimated separately for each model using Maximum A Posteriori (MAP) estimation with a multi-start optimization routine. Individual subject parameters were constrained using weakly informative priors (*α* ∼ Beta(2,5); *β* ∼ Gamma(2,3)). We performed model comparison using Bayesian Model Selection (BMS) at the random-effects level based on Bayesian Information Criterion approximations, evaluating expected posterior probabilities and protected exceedance probabilities to account for group heterogeneity

To validate the ability of our model comparison procedure to correctly identify the true underlying generative mechanism, we performed a model recovery analysis across all *Q*-learning models. For each generating model, we simulated 100 synthetic cohorts matching the empirical sample size (*n* = 38). True parameters (*α*, *β*) were drawn from uniform distributions (*α* ∈ [0.05,0.85], *β* ∈ [0.50,8.50]) to ensure the recovery procedure was not biased toward specific regions of the parameter space. Synthetic choices were generated using each model’s specific reward representation and task structure via a standard softmax choice rule. Each candidate model was subsequently fit to the identical simulated choice data using the same MAP optimization framework. Model recovery performance was quantified by constructing a confusion matrix of recovery proportions, reflecting how often each candidate model was selected as the best fit when data were generated by that model.

#### Self-report ratings

To assess the subjective experience elicited by individualized artificial stimuli, we compared ratings following BLA– and IT-targeting stimuli using paired *t*-tests (*n* = 39; separate tests were performed for valence and arousal). Differences in each of the seven emotion category ratings (happy, fear, sad, surprise, anger, disgust, and neutral) across synthesis targets (IT and BLA) were analyzed with a 2 × 7 repeated measures ANOVA (*n* = 39).

### Data availability

fMRI data, associated behavioral files, and experimental stimuli will be deposited to OpenNeuro and made publicly available as of the date of publication.

### Code availability

Code for all analyses is available at https://github.com/ecco-laboratory/AmygdalaManifold. Code for image synthesis is adapted from https://github.com/Animadversio/ActMax-Optimizer-Dev. Code for fMRI data analysis is available at https://github.com/canlab/canlabcore.

## Acknowledgements

This project was supported by grant R01MH134972 to PAK. The funders had no role in study design, data collection and analysis, decision to publish or preparation of the manuscript.

## Author Contributions Statement

Conceptualization, methodology, data collection, formal analysis, writing – original draft, writing – review and editing, validation, and visualization, GJ; data collection, formal analysis, writing – review and editing, and visualization, KR; data collection, writing – review and editing, EA; data collection, writing – review and editing, GT; conceptualization, methodology, formal analysis, writing – original draft, writing – review and editing, supervision, and resources, PAK.

