## Supplementary Materials for "A representation of the visual environment in the human basolateral amygdala supports learning and choice"

This PDF file includes:

Supplementary Figures 1-2
Supplementary Tables 1-8

### Supplementary Figures

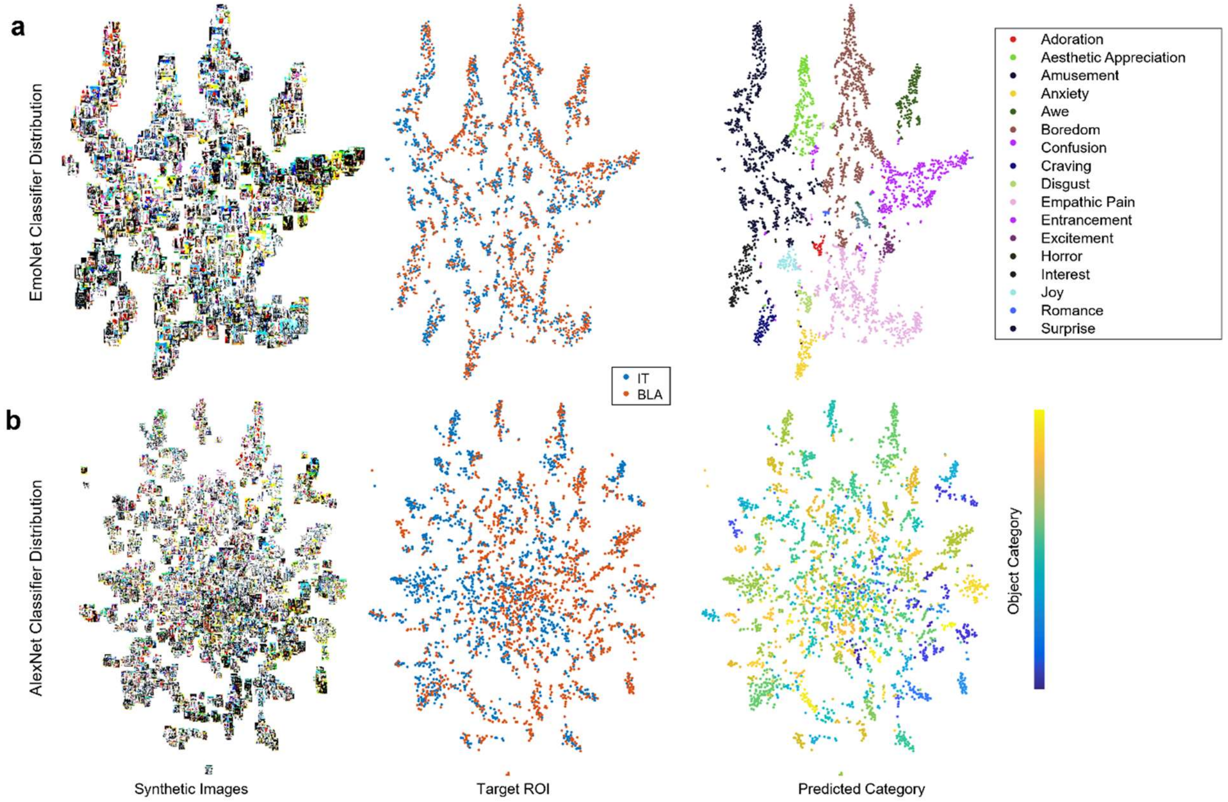


**Supplementary Figure 1. Feature organization of synthetic images targeting IT and BLA.** *t*-SNE projections of fc7 embeddings for images generated to target either the IT or BLA representation. **a**, Embeddings from the EmoNet model, shown for the full set of synthetic images (left), colored by the targeted ROI (middle), and colored by the predicted emotional category (right). **b**, Corresponding embeddings from the AlexNet model, shown for the full set of synthetic images (left), colored by targeted ROI (middle), and colored by the predicted object category (right). Each point represents a synthetic image. Emotion and object categories were predicted from the corresponding pretrained classifiers. The distributions provide a visualization of the high-level visual and affective features represented in the generated image set and their relationship to the targeted neural representations.

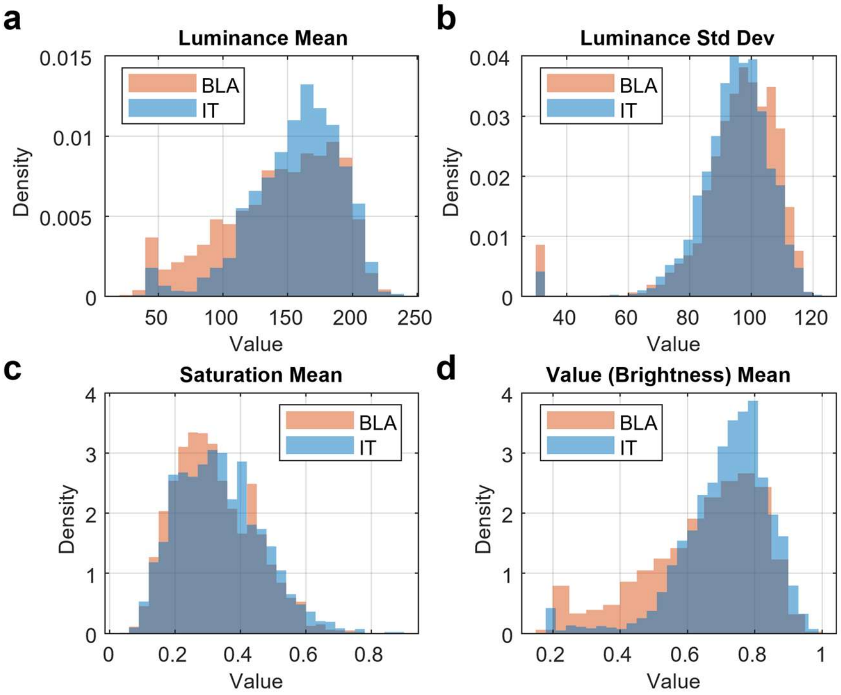


**Supplementary Figure 2. Comparison of low-level visual properties between BLA- and IT-optimized stimuli.** Probability density distributions for mean luminance, luminance standard deviation (spatial contrast), mean HSV saturation, and mean HSV value (brightness) calculated across all BLA-optimized (orange) and IT-optimized (blue) synthetic images. Both stimulus sets exhibit substantial overlap across all four parameters, demonstrating that optimization was not driven by systematic low-level visual differences.

#### Supplementary Tables

**Supplementary Table 1. Deep convolutional neural network predictions for amygdala optimized images**

| | EmoNet Classification | | | | | AlexNet Classification | | | | |
|---|---|---|---|---|---|---|---|---|---|---|
| Label | Boredom | Empathic Pain | Amusement | Entrancement | Aesthetic Appreciation | Pinwheel | Maraca | Comic Book | Ski | Admiral |
| Mean | 0.214 | 0.169 | 0.139 | 0.134 | 0.075 | 0.035 | 0.022 | 0.018 | 0.017 | 0.013 |
| SD | 0.318 | 0.293 | 0.260 | 0.268 | 0.199 | 0.097 | 0.099 | 0.058 | 0.062 | 0.081 |
| Cohen's *d* | 0.671 | 0.577 | 0.533 | 0.501 | 0.376 | 0.368 | 0.218 | 0.308 | 0.281 | 0.167 |

**Supplementary Table 2. Deep convolutional neural network predictions for inferior temporal cortex optimized images**

| | EmoNet Classification | | | | | AlexNet Classification | | | | |
|---|---|---|---|---|---|---|---|---|---|---|
| Label | Amusement | Boredom | Empathic Pain | Entrancement | Aesthetic Appreciation | Packet | Maraca | Park Bench | Mailbag | Balloon |
| Mean | 0.250 | 0.186 | 0.146 | 0.108 | 0.060 | 0.030 | 0.024 | 0.022 | 0.017 | 0.014 |
| SD | 0.342 | 0.304 | 0.258 | 0.225 | 0.183 | 0.079 | 0.078 | 0.102 | 0.054 | 0.089 |
| Cohen's *d* | 0.730 | 0.613 | 0.565 | 0.482 | 0.325 | 0.379 | 0.309 | 0.213 | 0.308 | 0.162 |

**Supplementary Table 3. ANOVA for self-reported emotion**

| Cases | Sum of Squares | *df* | Mean Square | *F* | *p* |
|---|---|---|---|---|---|
| Emotion[a] | 165,155.678 | 6 | 27,525.946 | 44.526 | < .001 |
| Residuals | 140,950.081 | 228 | 618.202 | | |
| ROI | 5.502 | 1 | 5.502 | 0.338 | .564 |
| Residuals | 618.665 | 38 | 16.281 | | |
| Emotion × ROI[a] | 160.854 | 6 | 26.809 | 1.126 | .348 |
| Residuals | 5,428.428 | 228 | 23.809 | | |

*Note.* Type III Sum of Squares

[a] Mauchly's test of sphericity indicates that the assumption of sphericity is violated ($p < .05$).

#### Supplementary Table 4. Mean differences for self-reported affect

*Paired Samples t-test*

| Measure 1 | | Measure 2 | *t* | *df* | *p* | Cohen's *d* |
|---|---|---|---|---|---|---|
| IT Arousal | - | BLA Arousal | 1.380 | 38 | .176 | 0.221 |
| IT Pleasantness | - | BLA Pleasantness | -0.948 | 38 | .349 | -0.152 |

*Note.* Student's *t*-test.

#### Supplementary Table 5. Reward values for each type of outcome image

| | BLA Image | IT Image | Blank Image |
|---|---|---|---|
| Approach Images | 1 | 1 | 0 |
| Avoid Blank Images | 0 | 0 | -1 |
| Approach BLA Images | 1 | 0 | 0 |
| Avoid BLA Images | -1 | 0 | 0 |
| Normative Amygdala | 1.478 (0.115) | 0.974 (0.093) | 0.563 (0.000) |
| Individual Amygdala | 1.856 (0.194) | 1.129 (0.160) | 0.262 (0.068) |
| Null | 0 | 0 | 0 |

#### Supplementary Table 6. Parameter Recovery

| Model | Correlation | | Bias | | Variance | |
|---|---|---|---|---|---|---|
| | α | β | α | β | α | β |
| Approach Images | .433 | .708 | -.188 | .021 | .006 | 3.581 |
| Avoid Blank Images | .542 | .640 | -.170 | .546 | .009 | 4.698 |
| Approach BLA Images | .374 | .639 | -.191 | -.248 | .005 | 3.031 |
| Avoid BLA Images | .519 | .626 | -.173 | .642 | .009 | 4.645 |
| Normative Amygdala | .485 | .706 | -.164 | -.071 | .013 | 4.233 |
| Individual Amygdala | .541 | .645 | -.167 | -.197 | .013 | 4.138 |
| Null | .023 | .021 | -.248 | -1.456 | .000 | .000 |

**Supplementary Table 7. Recovery of *Q*-Learning models**

| Simulated \ Predicted | Approach Images | Avoid Blank Images | Approach BLA Images | Avoid BLA Images | Population BLA | Individual BLA | Null |
|---|---|---|---|---|---|---|---|
| Approach Images | 1 | 0 | 0 | 0 | 0 | 0 | 0 |
| Avoid Blank Images | 0 | 1 | 0 | 0 | 0 | 0 | 0 |
| Approach BLA Images | 0 | 0 | 1 | 0 | 0 | 0 | 0 |
| Avoid BLA Images | 0 | 0 | 0 | 1 | 0 | 0 | 0 |
| Population BLA | 0 | 0 | 0 | 0 | 1 | 0 | 0 |
| Individual BLA | 0 | 0 | 0 | 0 | 0 | 1 | 0 |
| Null | 0 | .28 | .08 | .60 | .02 | .02 | 0 |

*Note.* Selection based on maximal BIC weight.

**Supplementary Table 8. Model comparison metrics for reinforcement learning models**

| | Approach Images | Avoid Blank Images | Approach BLA Images | Avoid BLA Images | Normative Amygdala | Individual Amygdala | Null |
|---|---|---|---|---|---|---|---|
| EPP | .0649 | .5317 | .0243 | .1969 | .0902 | .0399 | .0520 |
| EP | .0085 | .9506 | .0001 | .0360 | .0032 | .0007 | .0009 |
| BIC | 35.983 | 36.025 | 38.815 | 38.285 | 37.430 | 36.440 | 38.825 |
| ΔBIC | 2.955 | 3.000 | 5.787 | 5.256 | 4.402 | 3.411 | 5.796 |
| wBIC | .1456 | .2707 | .0824 | .1770 | .1196 | .1121 | .0926 |
